# Magnetic resonance imaging datasets with anatomical fiducials for quality control and registration

**DOI:** 10.1101/2022.11.21.516173

**Authors:** Alaa Taha, Greydon Gilmore, Mohamad Abbass, Jason Kai, Tristan Kuehn, John Demarco, Geetika Gupta, Chris Zajner, Daniel Cao, Ryan Chevalier, Abrar Ahmed, Ali Hadi, Bradley Karat, Olivia W. Stanley, Patrick Park, Kayla M. Ferko, Dimuthu Hemachandra, Reid Vassallo, Magdalena Jach, Arun Thurairajah, Sandy Wong, Mauricio C. Tenorio, Feyi Ogunsanya, Ali R. Khan, Jonathan C. Lau

## Abstract

Tools available for reproducible, quantitative assessment of brain correspondence have been limited. We previously validated the anatomical fiducial (AFID) placement protocol for point-based assessment of image registration with millimetric (mm) accuracy. In this data descriptor, we release curated AFID placements for some of the most commonly used structural magnetic resonance imaging templates and datasets. The release of our accurate placements allows for rapid quality control of image registration, teaching neuroanatomy, and clinical applications such as disease diagnosis and surgical targeting. We release placements on individual subjects from four datasets (n = 132 subjects for a total of 15,232 fiducials) and more than 10 brain templates (4,288 fiducials), compiling over 300 human rater hours of annotation. We also validate human rater accuracy of released placements to be within 1-2 mm (using a total of 50,336 Euclidean distances), consistent with prior studies. Our data is compliant with the Brain Imaging Data Structure (BIDS) allowing for facile incorporation into modern neuroimaging analysis pipelines. Data is accessible on GitHub (https://github.com/afids/afids-data).

## Background & Summary

Open resources available for reproducible, quantitative assessment of brain correspondence have been limited^1^. The most common metrics employed for the purpose of examining the quality of image registration, including the Jaccard similarity and Dice kappa coefficients, compute the voxel overlap between regions of interest (ROIs), which have been shown to be insufficiently sensitive when used in isolation or in combination for validating image registration strategies^1^. The ROIs used in voxel overlap are often larger subcortical structures that are readily visible on MRI scans (i.e., the thalamus, globus pallidus, and striatum), and thus lack the ability to detect subtle misregistration between images which may be crucial to detecting erroneous significant differences and variability^1-5^.

Inspired by classic stereotactic methods, our group created, curated, and validated a protocol for the placement of anatomical fiducials (AFIDs) on T1 weighted (T1w) structural magnetic resonance imaging (MRI) scans of the human brain^2^. The protocol involves the placement of 32 AFIDs found to have salient features that allow for accurate localization. The AFIDs are described using three-dimensional (x, y, and z) Cartesian coordinates and thus correspondence between points can be computed using Euclidean distances across a variety of applications. After a brief tutorial, AFIDs have been shown to have high reproducibility even when performed by individuals with no prior knowledge of medical images, neuroanatomy, or neuroimaging software. This was shown in separate studies where placements were performed on publicly available templates and datasets^2^ and a clinical neuroimaging dataset^3^.

The AFIDs protocol provides a metric that is independent of the registration itself while offering sensitivity to registration errors at the scale of millimeters (mm). This margin is crucial in neuroimaging applications (including morphometric analysis and surgical neuromodulation), where a few millimetres may represent the difference between optimal and suboptimal therapy.

The aim of this data descriptor is to provide the community with curated AFID placements and their associated MRI images. We release annotations on four datasets (n = 132; 15,152 fiducials) including healthy subjects and patients with neurological disorders, and more than 10 commonly used magnetic resonance imaging templates (4,288 fiducials), compiling more than 300 human rater hours of manual annotation of neuroanatomical structures. Descriptions of datasets and templates are provided in subsequent sections (see Table 1).

**Table 1:**
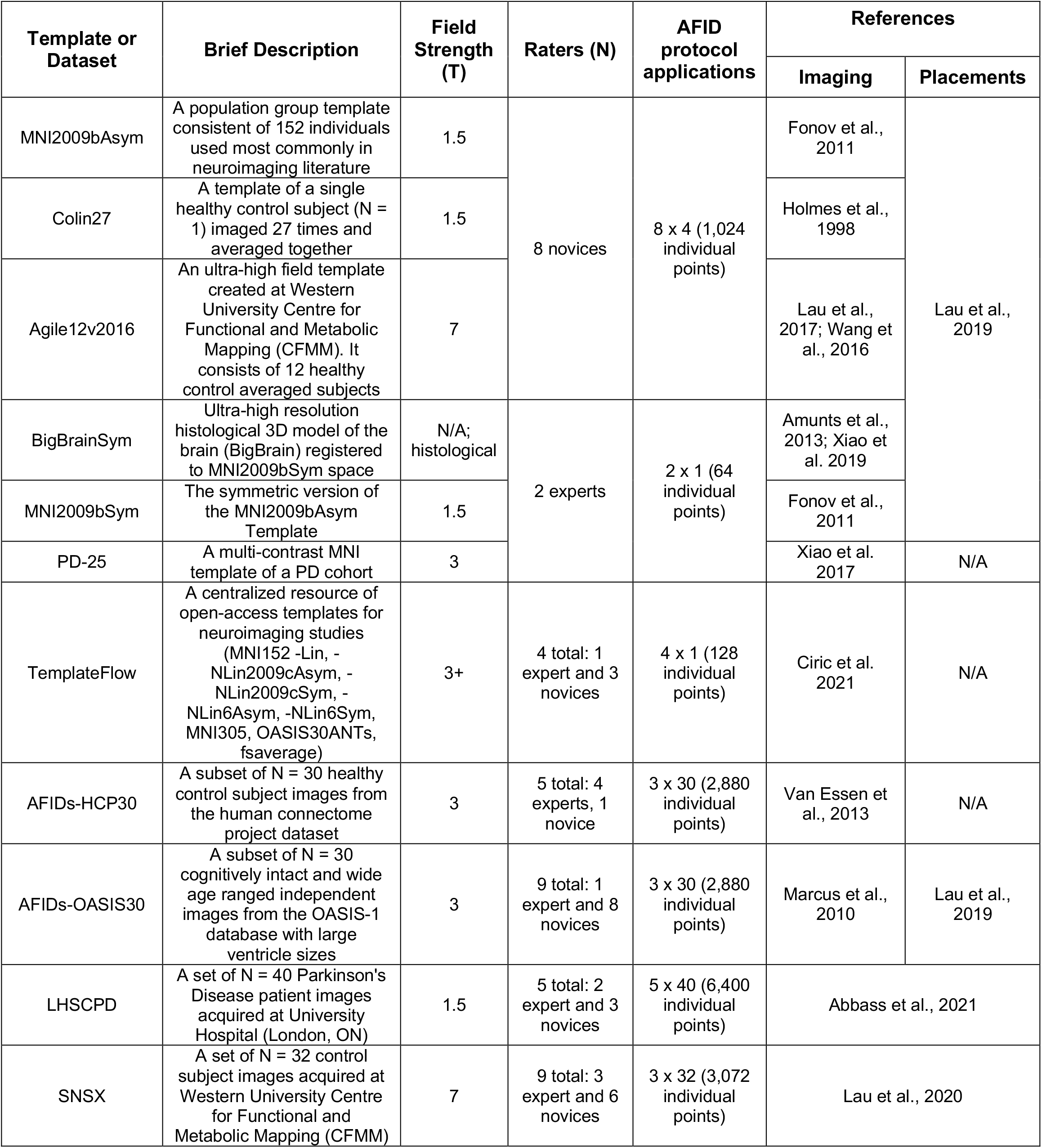
Summary of templates and datasets released, raters, and anatomical fiducial (AFID) protocol applications.

### Current Applications

#### Registration Assessment

We share our curated AFIDs annotations for a wide variety of datasets and templates of varying field strengths. This diversity of datasets will facilitate the testing and validation of image registration algorithms that can be used in many contexts. The user can select the datasets and templates that are in line with their neuroimaging application, then use the curated annotations to assess image registration quantitatively. For instance, AFIDs have been used to evaluate the process of iterative deformable template creation^6,7^, showing that error metrics generated from AFIDs converged differently as a function of template iterations and registration method (i.e., linear vs non-linear). Sharing the AFID placements and their associated images in the Brain Imaging Data Structure (BIDS) format aids in the convenience we strive to provide for the end-user and neuroimaging application developer^2,3,6,7^.

#### Education

New raters can compare their AFID placements to the curated normative distribution placements we release here. Our placements have been compiled over the years and can help raters assess accuracy for specific fiducials and subject/template data. To improve user accessibility and navigation of our released AFIDs annotations and framework, we also release the AFIDs validator (https://validator.afids.io). This tool provides: 1) detailed documentation of the AFIDs placement protocol, 2) an interactive way for users to upload placements to a regulated database, and 3) interactive ways to view uploaded placements relative to curated placements, which helps guide user to improve neuroanatomical understanding and placement accuracy^2,3^.

#### Brain structure and volumetric analyses

The 32 AFIDs (and associated images) in our pathologic dataset relative to the control can allow for insight on brain morphology and putative biomarkers of neurodegenerative diseases^3^.

### Prospective Applications

#### Registration optimization and quality control

The released imaging and AFID placement data may be useful in a few ways for improving neuroimaging pipelines: 1) providing centralized and quality controlled neuroimaging data (from more than 5 international neuroimaging datasets) allowing for a more accurate and generalizable head-to-head comparison amongst existing software for image registration, and 2) establishing a new registration metric which can be incorporated into neuroimaging software development workflows to optimize registration algorithm performance and also for quality control.

#### Automatic and accurate landmark placement

Our curated AFIDs can be used as ground truth placements when training machine-learning algorithms to automate brain landmark localization. Among the 32 AFID placements we release are the anterior and posterior commissures (AC and PC, respectively). Downstream applications of automatic localization include automatically computing AC-PC transforms (a common process in neuroimaging studies) and aspects of neurosurgical planning which involve the placement of these anatomical landmarks. The diversity of the released data (both hardware and disease status) will be crucial to the generalizability of such tools.

#### Surgical targeting

We release ultra-high field (7-Tesla; 7-T) MRI data where small structures like the subthalamic nucleus (STN)^8^ and zona incerta within the posterior subthalamic area are clearly visible^7^. Ground-truth locations of surgical targets (x, y, and z) can be related to the AFIDs placement locations via predictive models. This approach mitigates the lack of access to best case neuroimaging in clinical settings due to lack of high-field MRI or motion degradation.

#### Brain anatomy abstraction and anonymization

AFIDs and the distances between them represent an abstraction of brain anatomy in an anonymized way while still allowing for accurate pooling of data. Other major anatomical landmarks (representing lesions, tumors, or other structures) can be described in reference to the AFIDs “coordinate system” we establish using these curated placements.

## Methods

### Rationale for fiducial selection and placement assessments

The current version of the AFIDs protocol involves the placement of 32 anatomical fiducials. These AFIDs were selected to be easily identified on T1w MRI scans across varying field strengths (1.5-T, 3-T, 7-T) and were validated in previous studies^2,3^. During the selection process, regions that were prone to geometric inhomogeneity and distortion were avoided to enhance the accuracy of fiducial placement across applications of the AFIDs protocol^2^. There are 10 fiducials that fall on the midline and 11 located laterally on both hemispheres. The AFIDs protocol includes fiducials representing salient neuroanatomical features mostly located in the subcortex. Additional proposed fiducials could be included in future versions of the AFIDs protocol, but would require undergoing a similarly rigorous validation process^2,3^.

*Fiducial localization error (FLE)* is a term described by Fitzpatrick and colleagues^9^ that represents the distance between a fiducial position from its intended location. This term is used when operating image-guidance systems during neurosurgical procedures. In the context of the AFIDs protocol, and inspired by this extant terminology, we have defined the term anatomical fiducial localization error (AFLE). This value, in millimetres, can be thought of as the error arising from the placement (i.e., localization) of each of the 32 fiducials. When used to communicate the accuracy of all 32 AFIDs together, we term it global AFLE. There are three contexts for applying AFLEs: **1) Mean AFLE:** rater localization error relative to the intended location defined as the mean placement of all raters for a specific fiducial (termed ground truth AFID in subsequent sections). **2) Inter-Rater AFLE:** rater localization error calculated as the pairwise distances between different rater placements. If a single rater applied the AFIDs protocol more than once, then their mean placement coordinates were used for the pairwise distances calculations. **3) Intra-Rater AFLE:** rater localization error evaluating the precision of multiple placements by a single rater computed as the average pairwise distance between the same rater’s placements.

We also adopt the term *fiducial registration error (FRE)* in the context of the AFIDs protocol and term it the anatomical fiducial registration error (AFRE). It is important to note that FRE in our context diverges from the original usage by Fitzpatrick and colleagues^9^ which was restricted to external fiducials used in the context of image registration. Computed in millimetres, AFRE is defined as errors arising from the registration protocol applied on two images (often, but not limited to, subject and template). AFRE is the distance after co-registration between each of the 32 AFIDs placed on a subject image and their counterparts placed on template image. The average AFRE of all fiducials is termed the global AFRE. We also establish nomenclature to differentiate various use cases for AFRE. If an individual rater placement is chosen for subsequent analysis, then we term the resulting AFRE to be the **real-world AFRE** as it is more representative of what would happen in a clinical setting where one rater would apply the AFIDs protocol. If a ground truth AFID placement is used, then the resulting error is termed **consensus AFRE** as it represents the average placement among a group of raters prior to the image registration step. In this data descriptor, our focus is on releasing the curated fiducial placements and not an assessment of registration, so no AFRE metrics are produced. We still felt it would be useful to introduce AFRE as their computation constitutes one of the main applications of AFIDs and our shared datasets for quality control (i.e., in the context of image registration).

### Hardware and software used to curate data

All manually curated fiducials were placed using the Markups Module of 3DSlicer (an open-source imaging software)^10^. The datasets were curated at different times so a reference to the exact version of 3DSlicer and associated modules will be made under each dataset. 3DSlicer was chosen because it offers a variety of modules, particularly markups and registration modules were used for fiducial placement and AC-PC transform. 3DSlicer stores fiducials placed within its 3D coordinate system overlaid on the image giving the possibility of more accurate localization without the need to interpolate to the nearest voxel. The AFIDs placements released here for templates and datasets were performed on structural T1w MRI images.

### AFIDs protocol application

Before raters performed the AFIDs protocol, they attended a 3DSlicer workshop and placed the AFIDs protocol on a publicly available template as a form of training. For manual rater placements, the AFIDs protocol (https://afids.github.io/afids-protocol/) generally began with the placement of the anterior commissure (AC) and posterior commissure (PC) points (AFID01 and 02 respectively), which are defined to be at the center of each commissure. This was then followed by the identification of one or two more midline points (often the pontomesencephalic junction, AFID04, and the Genu of Corpus Callosum, AFID19, are used). After that, an AC-PC transform is performed, and the rest of the anatomical fiducials are placed. Rater placements deviating from a ground truth fiducial by greater than 10 mm were removed and considered outliers, as these errors are likely to be due to mislabelling and not reflective of true localization accuracy. In addition to subsequent sections, Table 1 provides brief descriptions of the released datasets and templates, information about raters, and AFIDs applications.

### AFIDs-HCP30 dataset

#### Subject demographics and imaging protocol

This subset consists of 30 unrelated healthy subjects (age: 21-52; 15 female and 15 male) chosen from the Human Connectome Project dataset (HCP). All scans were T1-weighted MR volumes with 1 mm voxels acquired on a 3-T scanner^11^.

#### Rater demographics and AFID placements

A total of 5 expert raters applied the AFIDs protocol. All raters had applied the AFIDs protocol before and have more than a year of neuroimaging, anatomy, and 3DSlicer experience. Three raters were previously heavily involved in validation studies^2,3^ and were assigned 10 random scans such that a total of one application of the AFIDs protocol was applied (via 3DSlicer 4.10.0). Two independent raters annotated all the 30 subjects for a total of three AFIDs protocol applications (2,880 fiducials) via 3Dslicer 4.10.0. Dataset can be found on: doi:10.18112/openneuro.ds004253.v1.0.3

### AFIDs-OASIS30 dataset

#### Subject demographics and imaging protocol

This subset consists of 30 subjects (age: 58.0 ± 17.9 years; range: 25-91; 17 female and 23 male) selected from the publicly available Open Access Series of Imaging Studies (OASIS-1) database^12^ and imaged at 3-T. The subjects were cognitively intact (Mini-Mental State Examination = 30), and the MRI scans were specifically chosen to be challenging (areas with more complex anatomy and asymmetries) by the senior author. More details on the selected subjects can be found in a previous study^2^. It is important to note that this subset of the OASIS-1 dataset is different from other currently existing subsets (for instance, the one used in the Mindboggle project^13^).

#### Rater demographics and AFID placements

Eight novice raters (11.5 ± 11.2 months imaging experience, 14.2 ± 17.0 months neuroanatomy experience, and 7.0 ± 8.8 months of 3DSlicer experience) and 1 expert rater (neurosurgical resident with 10 years experience in neuroanatomy) applied the AFIDs protocol via 3DSlicer 4.8.1. A total of 3 AFIDs protocol applications (2,880 fiducials) were performed as part of the AFIDs-OASIS30 dataset. Dataset can be found on: doi:10.18112/openneuro.ds004288.v1.0.2

### LHSCPD dataset

#### Subject demographics/template details and imaging protocol

The London Health Sciences Center Parkinson’s disease (LHSCPD) dataset currently consists of 40 subjects diagnosed with Parkinson’s Disease (age: 60.2 ± 6.8, range: 38 – 70; sex: 13 female and 27 male) with images acquired at University Hospital in London, ON, Canada on a 1.5-T scanner (Signa, General Electric, Milwaukee, Wisconsin, USA). The detailed imaging protocol was described in a previous study^3^. Ethics approval was received for anonymized release of patient scans by the Human Subject Research Ethics Board (HSREB) office at the University of Western Ontario (REB# 109045).

#### Rater demographics and AFID placements

There were 2 expert raters (over 5 years of experience in medical imaging, neuroanatomy, and 3DSlicer) and 3 novice raters (no knowledge of medical imaging, neuroanatomy, and 3DSlicer prior to training). AFIDs placements were performed using 3D Slicer version 4.10.0 on structural T1w images. A total of 5 AFIDs protocol applications were performed (6,400 fiducials). Dataset can be found on: doi:10.18112/openneuro.ds004298.v1.0.1

### SNSX dataset

#### Subject demographics and imaging protocol

The Stereotactic Neurosurgery (SNSX) dataset currently consists of 32 healthy participants (age: 46.2 ± 13.5 years; range: 20–70 years; 12 female and 20 male) with images acquired at the Western University Centre for Functional and Metabolic Mapping (CFMM) on a 7-T head-only scanner (Siemens Magnetom; Siemens Healthineers, Erlangen, Germany). An 8-channel parallel transmit/32-receive channel coil was used. The ethics approval, detailed imaging protocol, and pre-processing steps were documented in a previous study^7^.

#### Rater demographics and AFID placements

There were 3 expert and 6 novice raters recruited to apply the AFIDs protocol on the SNSX-32 dataset using 3DSlicer 4.8.1. No rater demographic data was collected, however, the 3 expert raters had more than 12 months of exposure to medical imaging, neuroanatomy, and 3DSlicer and applied the AFIDs protocol in our previous validation study^2^. The 6 novice raters had prior exposure to medical imaging, neuroanatomy, and 3DSlicer but have never applied the AFIDs protocol before training. The raters were split into 3 equal groups with one expert rater placed in each. Each group was randomly assigned a subset of the 32 subjects (two out of three rater groups had 11 subjects to annotate). Each rater within the group placed the AFIDs protocol on all subjects allocated to their group. Thus, the AFIDs protocol was performed a total of 3 times on all 32 subject scans (3,072 fiducials), with each rater annotating either 10 or 11 different subjects once depending on their group assignment. Dataset can be found on: doi:10.18112/openneuro.ds004241.v1.0.2

### MNI2009Asym & Agile12v2016 & Colin27 templates

#### Template details and imaging protocol

A group of commonly used public templates were annotated. The *MNI2009bAsym* is a population group template consisting of 152 individuals (aged 18.5–43.5 years) used commonly in the literature^14^. The images were acquired on a Philips 1.5-T Gyroscan (Best, Netherlands) scanner at the Montreal Neurological Institute.

The *Agile12v2016* is an ultra-high field template created locally at our institution (CFMM). It consists of 12 healthy control subjects (6 female; age: 27.6 ± 4.4 years). Scans were on a 7-T scanner (Agilent, Santa Clara, California, USA/Siemens, Erlangen, Germany) via a 24-channel transmit-receive head coil array^15,16^.

The *Colin27* is a template created from one subject scanned 27 times on a Phillips 1.5-T MR unit^17^.

#### Rater demographics and AFID placements

The same 8 novice raters that annotated the AFIDs-OASIS30 subset also annotated all of the templates mentioned above 4 times. Each rater performed the AFIDs protocol a total of 12 times for a total of 96 protocols (1,024 fiducials). Since raters annotated the same template more than once, there was an intra-rater metric calculated for these three templates (contrary to the datasets). Annotations were performed via 3DSlicer 4.8.1.

### BigBrainSym & MNI2009Sym & PD-25 templates

#### Template details and imaging protocol

BigBrain is an ultra-high resolution histological 3D model of the brain created using a large-scale microtome to cut a complete paraffin-embedded brain (65-year-old male) coronally at 20-mm thickness^18^. The BigBrainSym template refers to the BigBrain registered to MNI2009bSym space, defined in previous studies^2,6^. The MNI2009Sym is a symmetric version of the MNI2009Asym^14^.

The PD-25 template is a multi-contrast MNI template of a PD cohort with 3-T field strength^19^. We used the PD25-T1MPRAGE for the AFIDs placements.

#### Rater demographics and AFID placements

A total of 2 expert raters (more than one year of experience in neuroimaging, anatomy, and 3DSlicer and have been involved in prior validation studies^1,2^) were involved in placements. Each rater annotated both templates once (192 fiducials) via 3DSlicer 4.8.1.

### TemplateFlow templates

#### Template details and imaging protocol

All adult human structural MRI templates that could be found on TemplateFlow at the time of manuscript preparation were annotated (n=8)^20^.

#### Rater demographics and AFID placements

Three novice raters (no prior neuroimaging, anatomy, and 3DSlicer experience) and 1 expert rater (lead author; more than 10 years of experience in medical imaging, anatomy, and 3DSlicer) annotated a total of 8 templates (see Table 1). Each rater annotated the 8 templates once (1,024 fiducials) via 3DSlicer 4.10.

### AFLE calculation for all datasets and templates

All placements for a given scan and fiducial were averaged to achieve the ground truth fiducial placement per participant or template as shown in Figure 2a. For datasets, ground truth fiducial placements were computed for each subject in a dataset as shown in Figure 2b.

**Figure 1:**
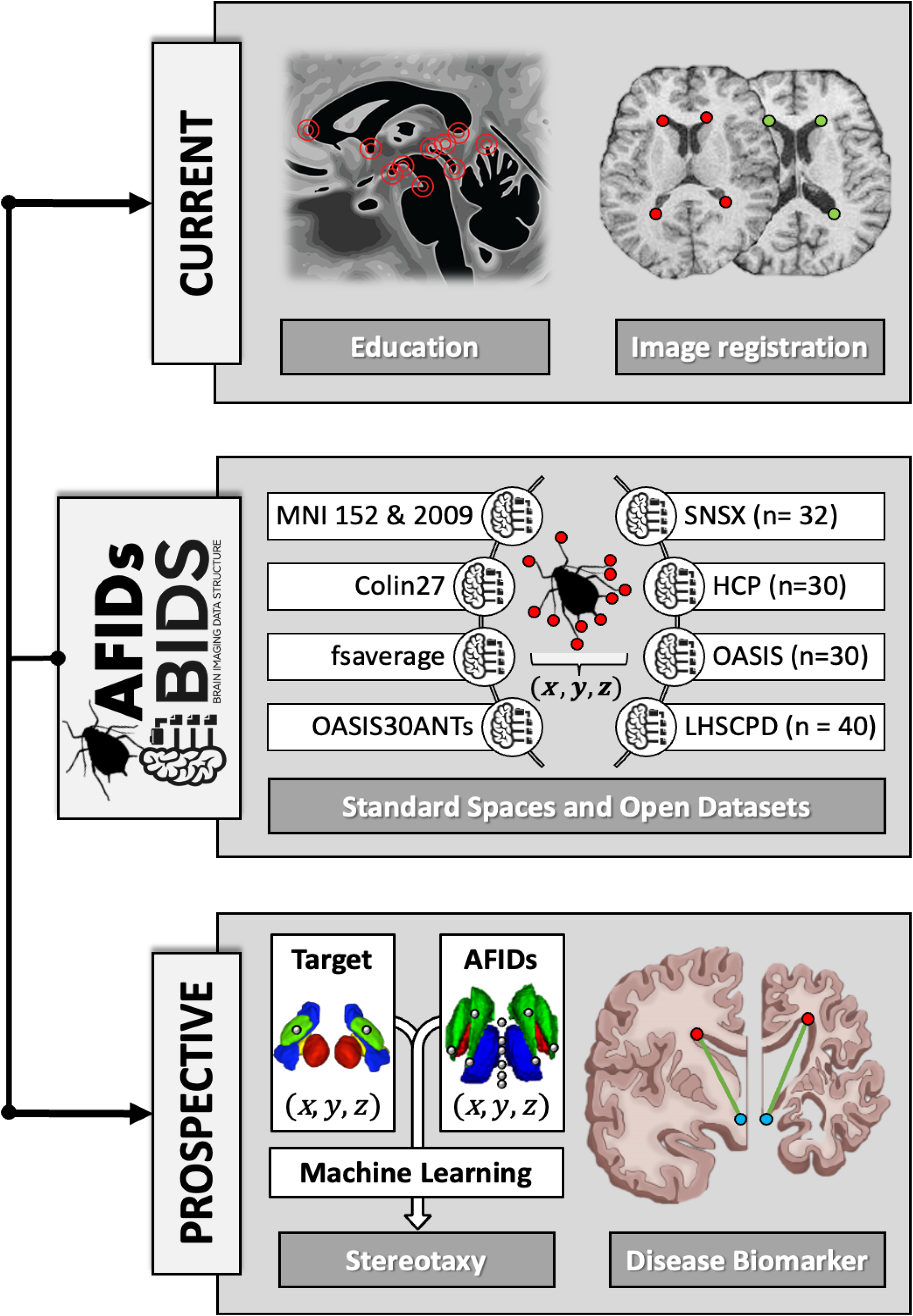
Current and prospective applications of curated anatomical fiducial (AFID) placements. Top panel: current applications in neuroanatomy education and image registration. Middle panel: released healthy and pathologic datasets and templates (descriptions can be found in text). Bottom panel: prospective applications of AFIDs in stereotactic targeting and as a disease biomarker.

**Figure 2:**
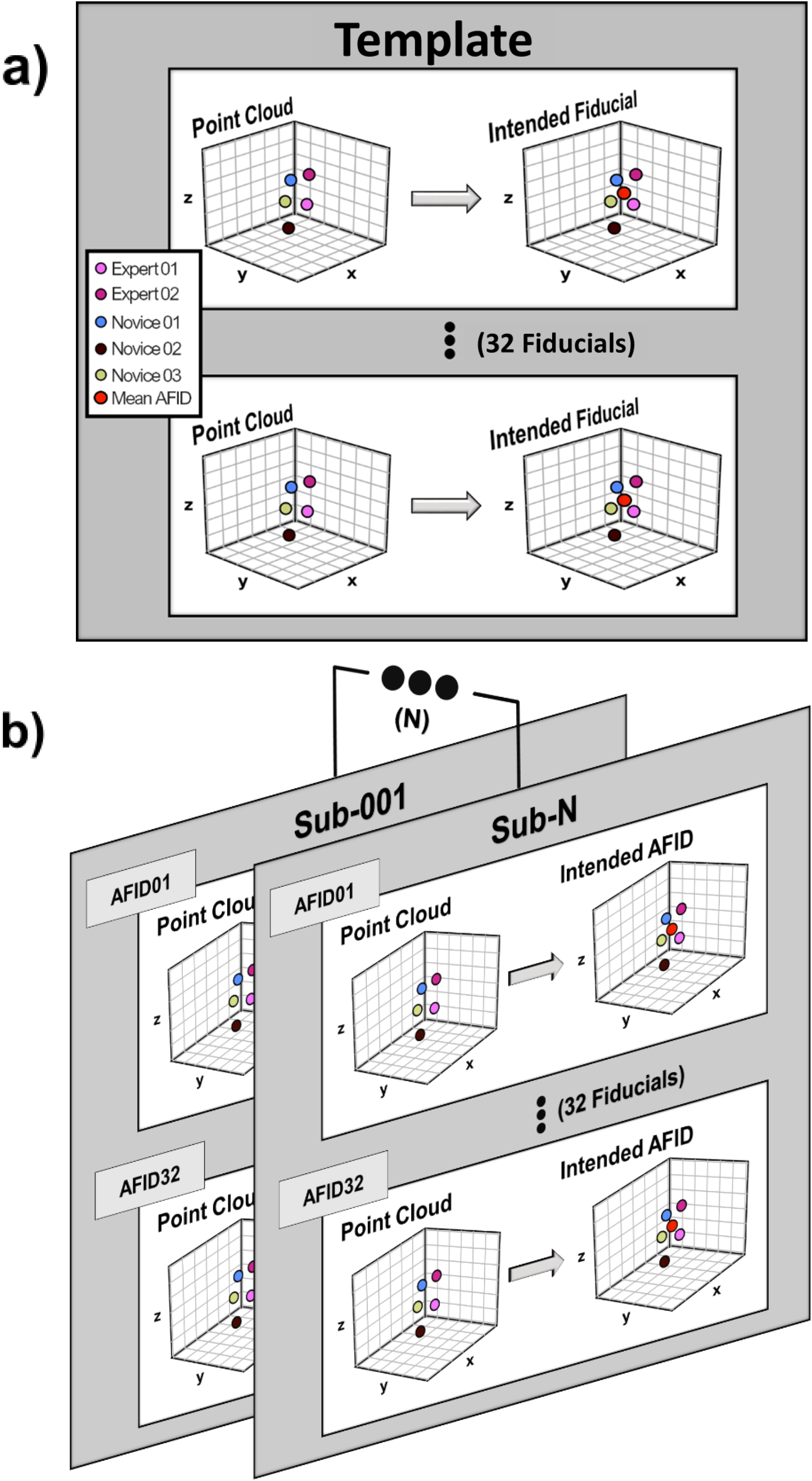
Ground-truth anatomical fiducial (AFID) placement on templates and datasets. (a) and (b) show the process of computing the intended AFID placement on a neuroimaging template or dataset respectively. It is the mean of the rater point cloud at each AFID, referred to as “ground-truth” in the text.

To compute the mean AFLE, Euclidean distances from the ground truth fiducial location to each of the individual rater placements were averaged for each fiducial. The result is termed the subject or template mean AFLE per fiducial. This process was independently repeated for all subjects. All subject mean AFLEs were averaged to achieve a dataset mean AFLE per fiducial as shown in Figure 3a. Finally, the dataset mean AFLE per fiducial was averaged across all fiducials to produce the global dataset mean AFLE. In a similar fashion, global inter-rater AFLE was computed for one subject across fiducials and then averaged across all subjects to produce a global dataset inter-Rater AFLE shown in Figure 3b.

**Figure 3:**
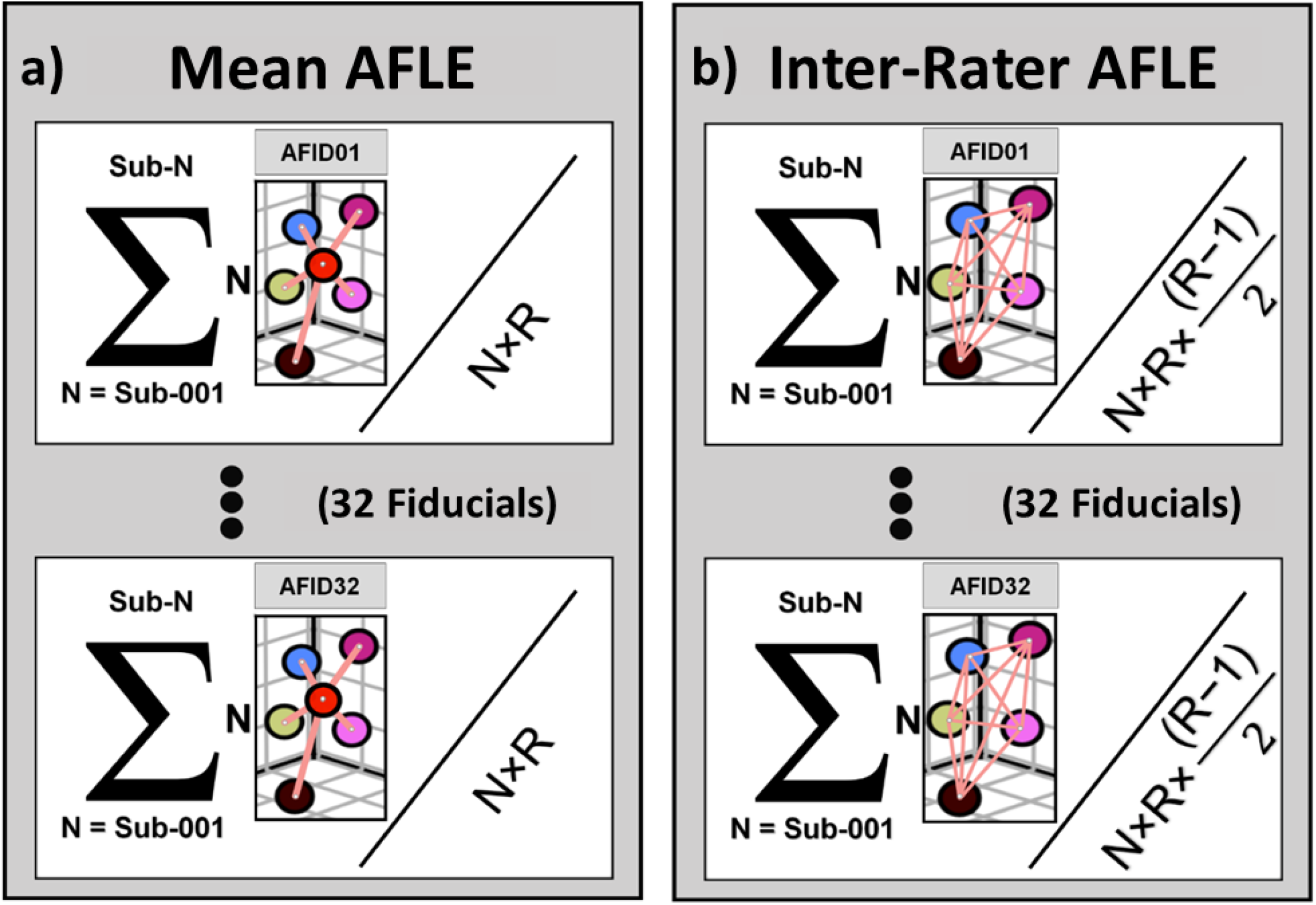
The technical validation computations for our anatomical fiducial (AFID) placements on templates and datasets. (a) and (b) show the equations used to compute mean and inter-rater anatomical localization error, respectively. *N = number of subjects in a dataset*. If calculating for a template, *N* would be 1. *R= the number of raters per scan/image*. In (a) Euclidean distances (shown in pink) represent distance from rater placement to the ground-truth (red). The mean AFLE was calculated by dividing the sum all Euclidean distances across all subjects (shown by the sigma notation) with the total number of Euclidean distances in the dataset (*N* × *R*) for each AFID. In (b) Euclidean distances (shown in pink) represent the pairwise distances between all rater placements on a scan. Inter-Rater AFLE was calculated by dividing the sum of the pairwise distances (shown by the sigma notation) by the total number of rater pairwise distances across a dataset per AFID 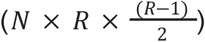.

## Data Records

In total, we release the curated AFID placements and associated imaging of 4 datasets and 14 openly available human brain templates (total of 19,520 manually placed anatomical landmarks — more than 300 human rater annotation hours). When available, individual rater placements were released, otherwise, the rater’s mean (ground-truth) placements were made available. The data we release here is BIDS compliant with a primary focus on adoption and usability. The AFIDs coordinates are described using the Markups comma-separated values file (i.e.,.fcsv extension) which is generated after the raters save their placements. The *.fcsv file is compatible for loading and viewing on 3DSlicer. As for the imaging data, all images used for annotations were BIDS compatible and made available in a compressed NIfTI-1 format (i.e.,.nii.gz extension). Each BIDS dataset has been released separately on OpenNeuro^21^ (links found under each dataset). A GitHub repository (serves as a centralized reference) also hosts AFIDs annotation files and template imaging data with directions for accessing BIDS datasets (https://github.com/afids/afids-data).

## Technical Validation

As mentioned in the methods, raters typically go through the AFIDs protocol by referring to the detailed documentation we have made available online (https://afids.github.io/afids-protocol/) and attend a neuroanatomy session with supplementary video (https://github.com/afids/afids-education). To ensure the placements we share are accurate and reproducible amongst expert and novice raters we computed the AFLE metrics to show the distribution in localization and validate that it is generally within 1-2 mm across various raters. Table 2 summarizes the AFLE metrics computed for each of the templates and datasets. On all datasets and templates, the mean AFLE metric was always within 1-2 mm.

**Table 2:** Summary anatomical localization errors (AFLE) and Euclidean distances (ED) used for their calculations for all released data

| Template or Dataset | EDs utilized for AFLE metrics | AFLE ± Error | |
| --- | --- | --- | --- |
|  |  | Mean | Inter-rater |
| MNI152NLin2009bAsym | Mean: 1,024 and inter-rater: 3,584 | $0.99 \pm 1.11$ | $1.07 \pm 0.46$ |
| Colin27 | | $1.71 \pm 2.78$ | $1.36 \pm 0.88$ |
| Agile12v2016 | | $1.10 \pm 1.59$ | $1.14 \pm 0.48$ |
| BigBrainSym | Mean: 64 and inter-rater: 32 | $0.63 \pm 0.50$ | $1.25 \pm 1.02$ |
| MNI152NLin2009bSym | | $0.55 \pm 0.26$ | $1.09 \pm 0.52$ |
| PD-25 | | $0.42 \pm 0.24$ | $0.83 \pm 0.47$ |
| MNI152Lin | Mean: 128 and inter-rater: 192 | $1.07 \pm 0.45$ | $1.74 \pm 0.74$ |
| MNI152NLin2009cAsym | | $1.03 \pm 0.40$ | $1.67 \pm 0.63$ |
| MNI152NLin2009cSym | | $1.06 \pm 0.47$ | $1.67 \pm 0.63$ |
| MNI152NLin6Asym | | $1.16 \pm 0.51$ | $1.90 \pm 0.86$ |
| MNI152NLin6Sym | | $1.08 \pm 0.54$ | $1.73 \pm 0.84$ |
| MNI305 | | $1.14 \pm 0.41$ | $1.85 \pm 0.52$ |
| OASIS30ANTs | | $0.78 \pm 0.33$ | $1.25 \pm 0.51$ |
| fsaverage | | $1.00 \pm 0.44$ | $1.65 \pm 0.73$ |
| AFIDs-HCP30 | Mean: 2,880 and inter-rater: 2,880 | $1.16 \pm 0.53$ | $1.98 \pm 2.03$ |
| AFIDs-OASIS30 | | $0.94 \pm 0.73$ | $1.58 \pm 1.02$ |
| LHSCPD | Mean: 6,400 and inter-rater: 12,480 | $1.57 \pm 1.16$ | $2.01 \pm 1.49$ |
| SNSX | Mean: 3,072 and inter-rater: 3,072 | $0.96 \pm 0.33$ | $1.64 \pm 1.37$ |
| **Total or Average** | **Mean: 19,520 and inter-rater: 30,816** | **$1.02 \pm 0.31$** | **$1.52 \pm 0.34$** |

## Usage Notes

We recommend loading the shared AFIDs annotation files (*.fcsv) in 3DSlicer alongside their associated images all of which are in BIDS format for ease of navigating. The local neuroimaging datasets we release here (namely, the LHSCPD and SNSX) will be quality controlled and expanded as more patients are recruited. Additionally, quality and version control of the AFIDs framework will be introduced as more collaborations and initiatives begin incorporating it into their workflows and releases. New templates and brain images can be added to future versions of the data descriptor once they have met standards for validation set by prior related studies^2,3^.

## Code Availability

GitHub repository for code used in technical validation and prior AFIDs studies can be found on the AFIDs project repository: https://github.com/afids/.

## Acknowledgements

The authors would like to acknowledge the efforts of all the students and trainees who created, maintained, and participated in neuroanatomy workshops and tutorials which established the foundations of the AFIDs framework. Additionally, all staff involved in patient recruitment, imaging, and maintaining locally curated datasets, namely the LHSC40PD and the SNSX-32 datasets.

Data was provided in part by Open-Access Imaging Series (Principal Investigators: D. Marcus, R, Buckner, J, Csernansky J. Morris; P50 AG05681, P01 AG03991, P01 AG026276, R01 AG021910, P20 MH071616, U24 RR021382)

Data was also provided in part by the Human Connectome Project, WU-Minn Consortium (Principal Investigators: David Van Essen and Kamil Ugurbil; 1U54MH091657) funded by the 16 NIH Institutes and Centers that support the NIH Blueprint for Neuroscience Research; and by the McDonnell Center for Systems Neuroscience at Washington University.

A.T. was supported by the Ontario Graduate Scholarship and Parkinson’s Southwestern Society Graduate Student Scholarship in partnership with the Mitacs accelerate program. A.K. was supported by the Canada Research Chairs program #950-231964, NSERC Discovery Grant #6639, and Canada Foundation for Innovation (CFI) John R. Evans Leaders Fund Project #37427, the Canada First Research Excellence Fund, and Brain Canada. J.L. was supported by research start-up funding through the Department of Clinical Neurological Sciences at University of Western Ontario.

## Author contributions

J.L., A.K., A.T., G.G. conceptualized the idea of the data release and obtained ethics approval. J.L., A.K., A.T., G.G., M.A., J.D., G.G., K.F. overlooked studies related to data release. A.T., J.L., and A.K. contributed to writing and preparing the manuscript. J.L., A.T., G.G., S.W., T.K., J.K., O.S. and M.A. provided and organized code for analysis and dataset organization. All authors contributed to data curation, material preparation, and reviewed the final version of the manuscript.

